# Environmental dynamics impact whether matching is optimal

**DOI:** 10.1101/2025.07.20.665805

**Authors:** Yipei Guo, Ann M Hermundstad

## Abstract

When foraging for resources, animals must often sample many options that yield reward with different probabilities. In such scenarios, many animals have been shown to exhibit “matching”, an empirical behavioral observation in which the fraction of rewarded samples is the same across all options. While previous work has shown that matching can be optimal in environments with diminishing returns, this condition is not sufficient to determine optimality. Furthermore, while diminishing returns naturally arise when resources in the environment deplete and take time to be replenished, the specific form of diminishing returns depends on the temporal structure and statistics of the replenishment process. Here, we explore how these environmental properties affect whether matching is optimal. By considering an agent that samples different options with fixed sampling rates, we derive the probability of collecting a reward as a function of these sampling rates for different types of environments, and we analytically determine the conditions under which the optimal sampling-rate policy exhibits matching. When all options are governed by the same replenishment dynamics, we find that optimality gives rise to matching across a wide range of environments. However, when these dynamics differ across options, the optimal policy can deviate from matching. In such cases, the rank-ordering of observed reward probabilities depends only on the qualitative nature of the replenishment process, but not on the specific replenishment rates. As a result, the optimal policy can exhibit under- or over-matching depending on how rewarding the different options are. We use this result to identify environmental settings under which performance differs substantially between matching and optimality. Finally, we show how fluctuations in these replenishment rates—which can represent either environmental stochasticity or the agent’s internal uncertainty about the environment—can accentuate deviations between optimality and matching. Together, these findings deepen our understand of the relationship between environmental variability and behavioral optimality, and they provide testable experimental predictions across a wide range of environmental settings.

## Introduction

To successfully forage for food and other resources, animals must contend with a wide range of environmental settings that vary in their resource availability. Given limited time and energy, a foraging animal must decide whether and when to explore these different settings in order to maximize its chance of success in collecting resources. In laboratory settings, this form of decision making is often studied by presenting animals with multiple simultaneously-available options that deliver reward—a proxy for an available resource in the environment [1]—with different frequency, and asking how animals allocate their choices to these different options [2–6]. Such experiments have shown that animals tend to exhibit “matching” behavior, an empirical observation that animals allocate their choices in proportion to the number of rewards that they collect from different options [7–14]. This phenomenon was initially observed in pigeons [7–11], but was subsequently observed in flies, mice, rats, monkeys, and humans [15–20]. In the original formulation of the “matching law”, if an animals makes *n*_*i*_ attempts at sampling an option *i*, and only *n*_*si*_ of those attempts are successful and yield reward, then matching is achieved when *n*_*i*_/Σ *n*_*i*_ = *n*_*si*_/Σ*n*_s*i*_ [7]. Equivalently, this implies that the fraction of successful attempts *n*_*si*_/*n*_*i*_ is the same across all options.

While data from early experiments in pigeons agree well with the matching law, many subsequent experiments revealed deviations from precise matching. In particular, animals often have a tendency to sample better options less frequently than one would expect from precise matching, a phenomenon known as “under-matching” [14,19,21]. Given the prevalence of these different experimental observations, it is natural to ask why matching is so commonly observed, whether it is a feature of optimal behavior, and under what conditions we should expect strict matching versus deviations from it.

Many previous studies have explored the relationship between matching and optimality in the context of specific experimental setups and task structures. These include the commonly-used concurrent variable-ratio schedule, where a reward is available at an option after a random number of attempts at that option, and the concurrent variable-interval schedule, where a reward is available after a random amount of time has passed [2–6]. Recent work has considered matching within a more general context, where it has been tied to optimality through diminishing returns [22]. However, diminishing returns are not sufficient to guarantee that optimality gives matching [22], and they can manifest in different ways depending on how resources are depleted and replenished in the environment.

In this work, we explore how different environmental dynamics affect whether the optimal policy gives rise to matching. We model an environment that produces and replenishes resources over time. These resources can be collected at different sites (“options”), that can be governed by different replenishment dynamics. We decompose these dynamics into separate contributions that control the structure, statistics, and overall quality of the replenishment process, and we ask how an agent should best allocate its choices to exploit these properties and maximize the number of resources that it can collect. To this end, we consider a simple, memoryless agent that samples each available option at a fixed rate, subject to a maximum total sampling rate. We then derive conditions under which the optimal policy that maximizes resource collection across options will also exhibit matching. We show that while the optimal policy gives rise to matching in many types of environments, this may not be the case when the nature of the replenishment process differs across the options. In such cases, we show how the observed deviations from matching depend on the quality and reliability of the environment. In doing so, we provide concrete predictions about the impact of different environmental manipulations on both optimal and matching behavior.

## Results

We consider an environment that contains *N* distinct options, each of which delivers discrete resources that replenish over time. At any given time, each option can be in one of two states: empty (contains a resource) or full (does not contain a resource). An agent can collect a resource by sampling an option that is in the full state; the option then immediately depletes to the empty state (Fig. 1a). A replenishment process governs the time at which an empty option reverts back to the full state. The success of the agent in collecting resources depends on the agent’s behavioral policy, which determines how the agent samples different options over time, and on the dynamics of the replenishment process, which governs when resources become available at different options. Below, we first characterize these two components and then study their interaction.

**Figure 1.**
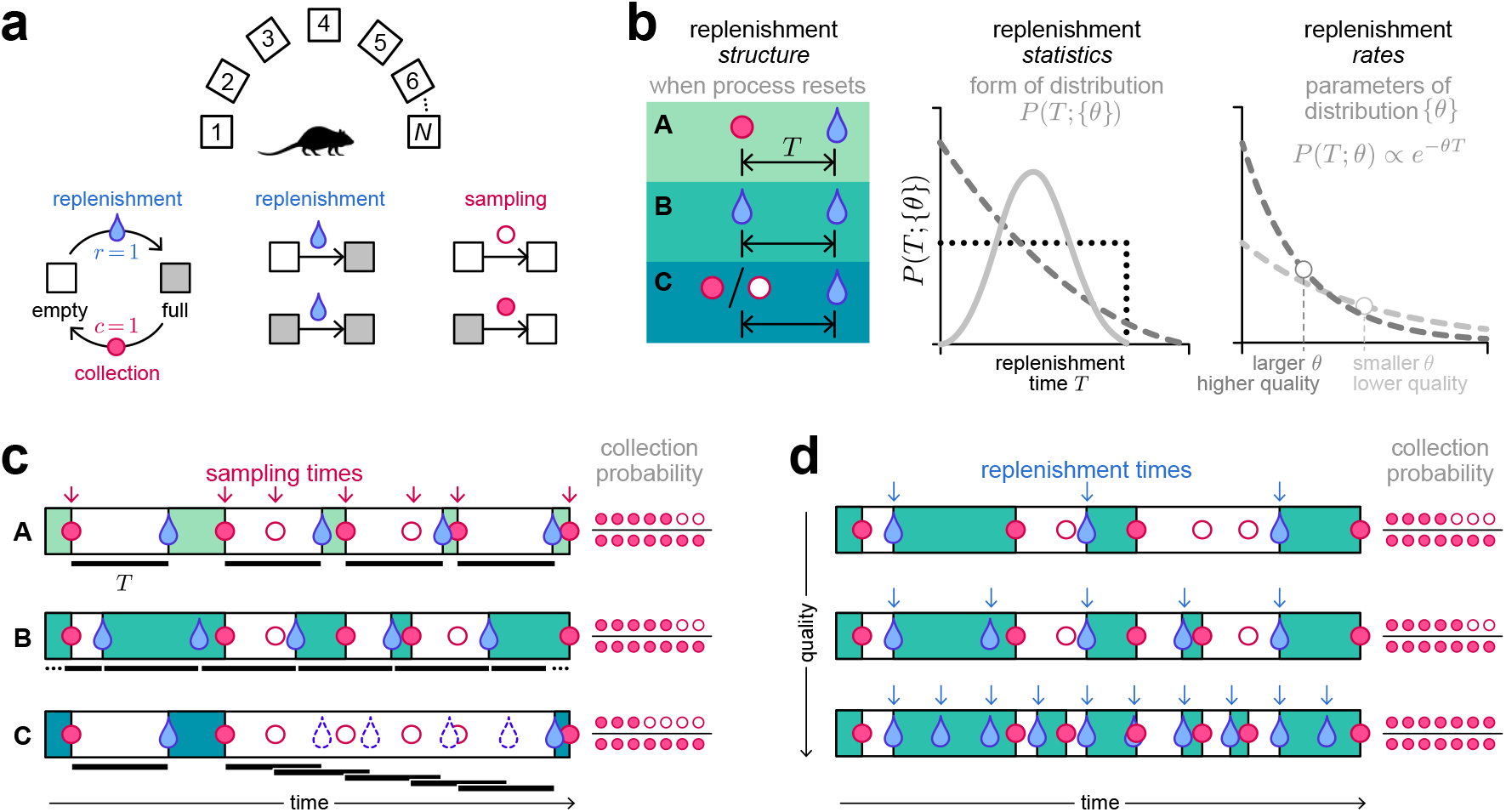
Depleting options can be characterized in terms of the structure and statistics of replenishment. **(a)** We consider an environment with *N* options that deliver discrete resources and that replenish after resources are collected. Each option can be in one of two states: empty or full. An option is always in the full state after a replenishment event. After a sampling event, the option transitions to the empty state. If an option is full when sampled, the sampling event was successful and the resource is collected; if an option is empty when sampled, the sampling event was unsuccessful and nothing is collected. **(b)** We differentiate three distinct features of the replenishment process: (i) the replenishment structure, which refers to *when* the process resets and specifies the meaning of the replenishment time *T*(left); (ii) the replenishment statistics, which refers to the form of the distribution *P* (*T*;{θ}) (middle); and (iii) the replenishment rates, which refer to the parameters {θ} of that distribution (right). The first two features control qualitative features of the replenishment process; the third feature quantitatively determines the overall quality of the environment, with higher replenishment rates parameterizing higher quality environments that replenish more quickly. **(c)** Each replenishment structure can yield different numbers of collections, even for the same sampling times (red arrows and circles). **(d)** Higher quality environments replenish more quickly (blue arrows), and thus yield more collections for the same sampling times (red circles). Illustrated for replenishment structure B.

### Dynamics of replenishment

We characterize three distinct features of the replenishment process (Fig. 1b): (i) the *structure* of replenishment, which controls *when* the process resets by specifying how the replenishment time *T*is measured relative to the agent’s choices or outcomes; (ii) the *statistics* of replenishment, which specifies the form of the distribution *P*(*T*; {*θ*}) of replenishment times; and (iii) the *rates* of replenishment, which specify the parameters {*θ*} of the distribution *P*(*T*) and control the overall quality of the environment (i.e., whether resources replenish quickly or slowly).

We consider three distinct types of replenishment structures (Fig. 1c):

A. The process resets after each resource collection (Fig. 1c, top row). Imagine the scenario where a worker always keeps an eye on an item on a shelf in a grocery store, and only restocks the item whenever it runs out. The replenishment process (which involves the worker going to the storage room, retrieving the item, and bringing it to the shelf) resets whenever the item is depleted from the shelf, and *T*is measured from the most recent collection event. In this case, any sampling attempts made between the last collection and the next replenishment event would be unsuccessful. This corresponds to the scenario implemented under the interval schedules that are commonly used to study decision-making in animals [23, 24].
B. The process resets independently of the agent’s actions or the state of the option, in which case *T*is measured from the most recent replenishment event (Fig. 1c, middle row). For example, suppose we were to leave a small pail out in the open to catch water whenever it rains. Whether we choose to sample the pail (i.e., check on the pail and use up any available water) does not affect when it is next going to rain.
C. The process resets after each sampling attempt, regardless of whether a resource was collected (Fig. 1c, bottom row). In natural environments, this scenario could arise in settings where the agent’s actions disrupt the replenishment process. For example, suppose one wishes to collect honey from a beehive. The production of honey requires a large population of bees, but the sampling process scares them away. After every sampling event, it takes some time for the bees to return and start producing honey. In this case, the replenishment process resets whenever sampling occurs, regardless of whether honey is collected, and *T*is measured from the most recent sampling event. A key feature of this type of replenishment structure is that a higher sampling rate does not necessarily lead to a higher number of collections, because over-sampling can result in repeated disruptions of the replenishment process. Given the same distribution *P*(*T*) and the same sampling times, this structure results in more unsuccessful attempts than structures (A) and (B) (Fig. 1c).

Within each of these structures, the replenishment times are fully specified by the distribution *P*(*T*; {*θ*}). Higher replenishment rates *θ* give rise to higher quality environments in which the agent collects more resources under the same sampling times (Fig. 1d).

### Behavioral policy

The success of the agent in collecting resources depends on how its behavioral policy interacts with the replenishment processes described above. To characterize this interaction, we focus on a class of “fixed-sampling-rate” policies that specify the frequency with which the agent samples each option. This class of policies has been studied previously [22] and enables us to analytically derive conditions under which optimal behavior exhibits matching.

Within this class, the agent’s behavioral policy is fully specified by a fixed vector of sampling rates 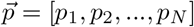. We assume that the agent operates under a constrained energetic budget, such that the overall sampling rate cannot exceed a maximum value: Σ_*i*_ *p*_*i*_ ≤1 (Fig. 2a). This constrains the long-term average number of sampling attempts per unit time. These rates govern when and which option the agent samples relative to the underlying dynamics of the replenishment processes, and thus determine whether or not the agent will collect a resource from any given sampling event. Fig. 2b illustrates this for a single option that replenishes after a fixed time interval. If the agent samples infrequently, it is guaranteed to collect a resource. However, as the agent increases its sampling rate, a larger fraction of its samples are unsuccessful because the option has been depleted and has not yet replenished. As a result, the average probability that a given sampling event results in a successful collection— which we refer to as the collection probability *P*_*ci*_(*p*_*i*_)—decreases with the sampling rate *p*_*i*_ (Fig. 2c, top). The *rate* at which resources are collected, *c*_*i*_ = *p*_*i*_*P*_*ci*_(*p*_*i*_), depends on both *p*_*i*_ and *P*_*ci*_(*p*_*i*_) and can therefore increase or decrease with *p*_*i*_ depending on how *P*_*ci*_(*p*_*i*_) decays with *p*_*i*_ (Fig. 2c, middle). However, the collection rate always initially increases with sampling rate, and the marginal gain *g*_*i*_(*p*_*i*_) = *dc*_*i*_/*dp*_*i*_ always initially decreases with sampling rate (Fig. 2c, middle and bottom). Both of these properties are characteristic of diminishing returns; more formally, an option that exhibits diminishing returns will satisfy 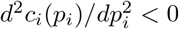.

**Figure 2.**
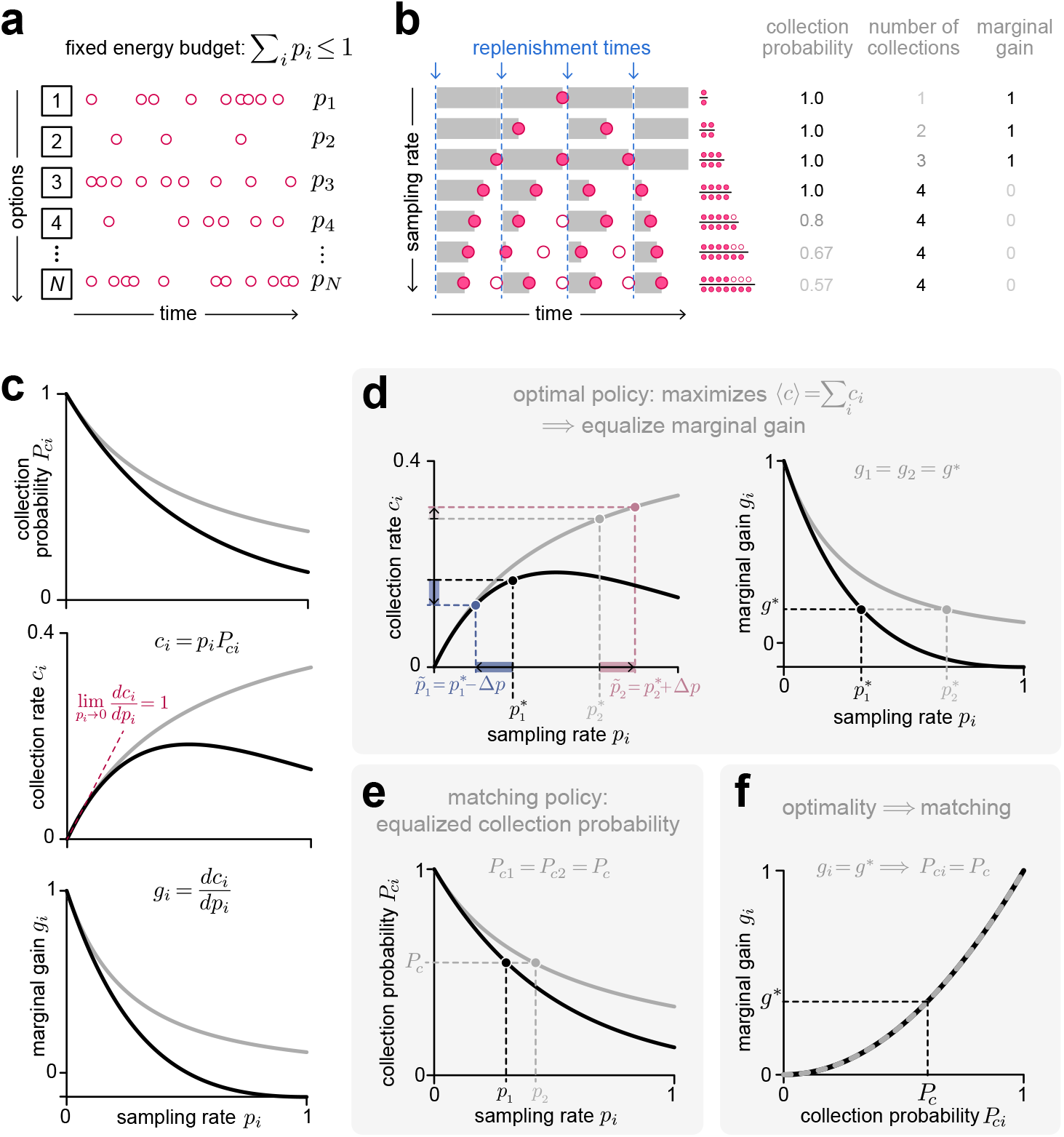
Sampling rates interact with the replenishment process to determine how many resources the agent collects. (**a)** We assume that the agent chooses each of the *N* options with a fixed sampling rate *p*_*i*_, subject to a constrained energetic budget Σ_*i*_ *p*_*i*_ ≤ 1. **(b)** Illustration of a simple environment in which a single option replenishes at fixed time intervals. Increasing the sampling rate leads to a reduction in the collection probability, an increase in the total number of collections, and a decrease in the marginal gain (i.e., the increase in number of collections for an increase in sampling rate). **(c)** Features of collection as a function of sampling rate, shown for two different options that differ in their replenishment dynamics but both exhibit diminishing returns. Top: because it takes time for resources to replenish, the collection probability *P*_*ci*_ at a given option *i* monotonically decreases with sampling rate *p*_*i*_. Middle: the collection rate *c*_*i*_, which has contributions from both *p*_*i*_ and *P*_*ci*_ (and can therefore increase or decrease with *p*_*i*_), initially increases with *p*_*i*_ at the same rate for all options. Bottom: the rate at which *c*_*i*_ increases with *p*_*i*_—which we refer to as the marginal gain *g*_*i*_—decreases as *p*_*i*_ increases. **(d)** Under the optimal policy that maximizes the net collection rate, the marginal gain is equalized across options. Left: for the two options illustrated in panel **c**, the optimal sampling rates 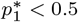 and 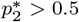 yield the same marginal gain across options (i.e.,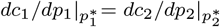). Altering the sampling rates away from these values, for example by increasing 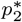 and decreasing 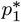, would lead to an increase in the collection rate at option 2 (red), but would be outweighed by a larger decrease in the collection rate at option 1 (blue). Right: As a result of the properties illustrated in the left panel, the optimal policy 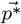 that maximizes the net collection rate must satisfy Eq. 1, such that the marginal gain is equalized across options (i.e., *g*_1_ = *g*_2_ = *g*^*∗*^). **(e)** A policy exhibits matching if the collection probabilities are equalized across options (i.e., *P*_*c*1_ = *P*_*c*2_ = *P*_*c*_). **(f)** Optimality gives matching if collection probabilities are equalized across options under the optimal policy, such that the optimal sampling rates that satisfy *g*_*i*_ = *g*^*∗*^ also satisfy *P*_*ci*_ = *P*_*c*_. This is always satisfied when the function *g*(*P*_*c*_) is the same across options.

We define the optimal allocation of sampling rates to be the one that maximizes the net collection rate across all options,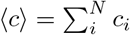. This optimal policy 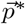 satisfies the condition 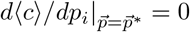 for all options *i*; in other words, any changes in sampling rates will decrease the net collection rate. Since the initial marginal gain 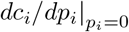 takes on the same value for all possible replenishment dynamics that we consider here, this condition implies that the marginal gain should be equalized across options (SI Section 1) [22]:

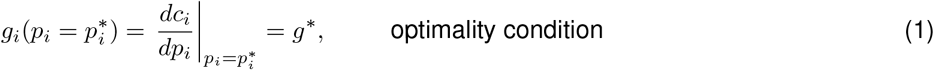

where *g*^∗^ is the lowest non-negative value that can be achieved without exceeding the total sampling rate budget.

This optimality condition can be intuitively understood by considering the two different options shown in Fig. 2c. The first of these options, in black, exhibits a more rapid decay in collection probability with sampling rate. As a result, the marginal gains of both options can be equalized if the agent samples the second option more frequently than the first (Fig. 2d); for a fixed maximal energy budget *p*_1_ + *p*_2_ = 1, this is achieved when *p*_1_ < 1/2 and *p*_2_ > 1/2. To maintain this energy budget, an increase in one sampling rate (e.g. *p*_2_) must be offset by a decrease in the other (*p*_1_). This will necessarily lead to a decrease in the net collection rate, because the small increase in collection rate from increasing *p*_2_ will be offset by a larger decrease in collection rate from decreasing *p*_1_. As a result, any deviation away from this policy that produces equalized marginal gains will lead to an overall reduction in the net collection rate, implying that this policy is optimal.

In comparison, if a policy gives rise to matching, the collection probabilities will be equalized across options (Fig. 2e):

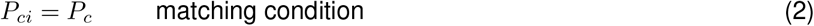

Thus, optimality gives rise to matching if collection probabilities are matched under the optimal policy (i.e., if both the optimality and matching conditions are satisfied).

In any given environment, the optimal policy—and the corresponding marginal gain *g*^∗^ and collection probabilities 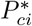– will depend on the features of replenishment at each option (Fig. 1b). If the relationship *g*(*P*_*c*_) between the marginal gain and the collection probability is the same across options, optimality will always give rise to matching, since any policy that equalizes marginal gains (including the optimal policy) will also equalize collection probabilities (Fig. 2f). However, in general, *g*(*P*_*c*_) may not be the same across all options. In what follows, we derive *g*(*P*_*c*_) for different types of environments, and we use it to study how the properties of replenishment affect whether optimality gives rise to matching.

### Optimality gives matching when all options share the same qualitative features of replenishment

To determine whether the optimal policy exhibits matching for any given environment, we first derive the expression for the collection probability *P*_*ci*_(*p*_*i*_) given a policy of fixed sampling rates *p*_*i*_. We then use this to compute the collection rate *c*_*i*_(*p*_*i*_) = *p*_*i*_*P*_*ci*_(*p*_*i*_) and marginal gain 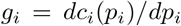 for each option. In principle, each of these quantities can depend on the specific form and rate of replenishment at each option, and there is no guarantee that a given policy will generically equalize both the collection probability and the marginal gain. However, we will show that in some settings, both the collection probability and marginal gain can be written as functions of the policy alone, without additional dependence on the form or rates of replenishment. In such cases, the marginal gain can be expressed as a function of the collection probability (i.e., *g* ≡ *g*(*P*_*c*_)), such that optimality always gives matching.

#### If all options replenish after resource collection, the optimal policy exhibits matching

Because resources take time to replenish, any sampling attempt made between a resource collection and a replenishment event will be unsuccessful (all other attempts are, by definition, successful). When the replenishment process is triggered by the collection itself (replenishment structure A), the number of unsuccessful attempts *n*_*u*_(*T*) is a function of the replenishment time *T*. The steady-state collection probability when sampling an option *i* can thus be written as (Fig. 3a):

**Figure 3.**
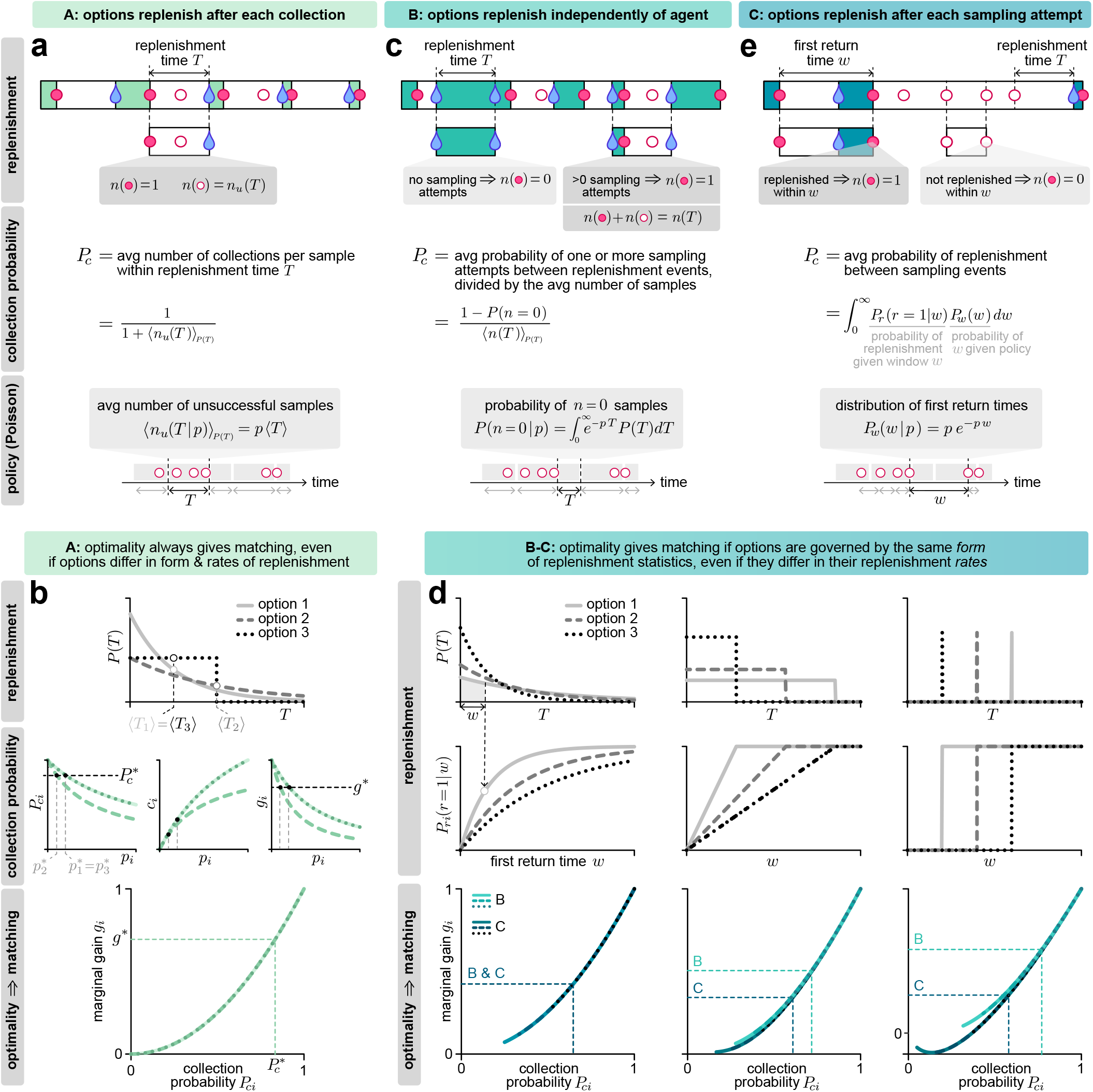
The optimal policy exhibits matching in many settings. **(a)** When options replenish after each collection, the collection probability can be written as the average number of collections per samples within a replenishment time T. For a fixed sampling-rate *p*, this depends only on the average replenishment time ⟨T⟩ and not on the function form of *P* (T). **(b)** Consider three options that differ in the form and rate of replenishment (upper). The collection probability, collection rate, and marginal gain are identical for the two options that share the same rate of replenishment, even though they are governed by different forms of replenishment statistics *P* (*T*) (middle). All three options share the same relationship between the marginal gain and the collection probability, such that matching is always optimal (lower). **(c)** When options replenish independently of the agent, the collection probability can be written as the average probability of sampling at least once between replenishment events, divided by the average number of samples between replenishment events. **(d)** Consider three options that share the same form of replenishment statistics, but differ in their rates (top and middle rows). As long as the options are governed by the same replenishment structure (shown here for structures B and C), they share the same relationship between the marginal gain and the collection probability, and hence, optimality always gives rise to matching. Note that when plotting relationships between marginal gain and collection probability, we accounted for the maximum sampling rate constraint and hence only included regions where *p*_*i*_ ≤1. **(e)** When options replenish after each sampling attempt, the collection probability can be written as the average probability that a replenishment will fall between two sampling attempts, which depends on the distribution of first return times.

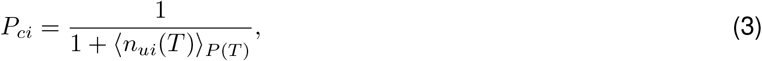

where ⟨·⟩_*P* (T)_ indicates an average over the distribution of replenishment times.

In general, *n*_*ui*_(*T*) is dependent on the form of the policy. However, for the fixed sampling rate policy that we consider here, *n*_*ui*_(*T*) = *p*_*i*_*T*. Therefore, Eq. 3 can be written as (Fig. 3a):

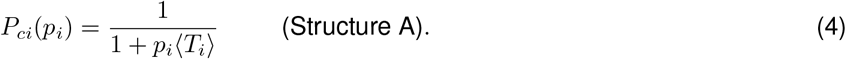

With this simplification, it is straightforward to show that when options replenish after resource collection, the optimal policy always exhibits matching. To see this, we can use the expression for *P*_*ci*_(*p*_*i*_) to write the collection rate 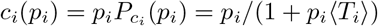, which can be used to calculate the marginal gain:

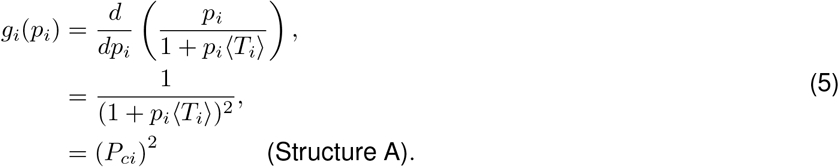

Because *g*_*i*_ depends only on *P*_*ci*_, the optimal policy (that satisfies *g*_*i*_ = *g*_j_) will also give rise to matching (*P*_*ci*_ = *P*_*c*j_). This is true regardless of the values of ⟨T_*i*_⟩ and the form of *P*_*i*_(*T*), and even if the form of *P*_*i*_(*T*) differs across options (Fig. 3b).

#### If all options are governed by the same replenishment process, the optimal policy exhibits matching

When the replenishment process is triggered by events other than collections, it is not straightforward to calculate the distribution of times between a collection and the next replenishment. Instead, we use other properties of the replenishment process to directly compute the collection probability.

##### Options replenish independently of the agent

When the replenishment process resets independently of the agent’s choices (replenishment structure B), only the first of the *n*_*i*_ sampling attempts that fall between two consecutive replenishment events will be successful; all other attempts will be unsuccessful. The average probability of collecting a resource within an interval *T*is thus the average probability that there is at least one sampling attempt in that interval, divided by the average number of sampling attempts in that same interval (Fig. 3c):

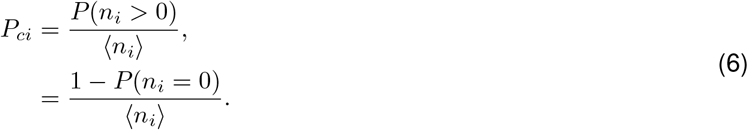

This derivation does not rely on any properties of the behavioral policy, and thus holds for any arbitrary policy type. For the fixed sampling rate policy we consider here, the average number of attempts that fall between two consecutive replenishment events is ⟨*n*_*i*_⟩ = *p*_*i*_ ⟨T_*i*_⟩ = *p*_*i*_ *⎰ TP*_*i*_*(T)dT* and the average probability that there are no sampling attempts within that same interval is *P*(*n*_*i*_ = 0) = ⎰ exp(−*p*_*i*_*T*)*P*_*i*_(*T*)d*T*. Moreover, if all options have the same form of replenishment statistics and differ only in their replenishment rates, we can write *P*_*i*_(*T*) = *P*(T|θ_*i*_), where θ_*i*_ is a parameter that controls the replenishment rate of each option. After defining the re-scaled variables 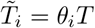 and 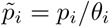, the average collection probability can be written as:

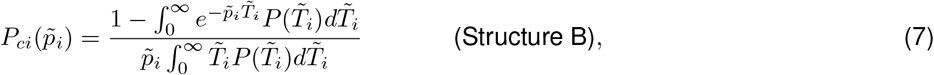

and the corresponding marginal gain is given by:

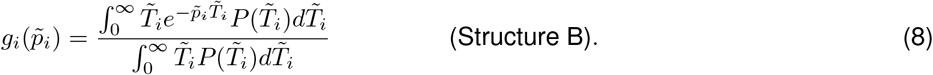

Because both functions depend solely on 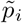, they are both equalized when 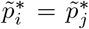 and thus opyimal policy again exhibits matching (Fig. 3d).

##### Options replenish after each attempt

When the replenishment process resets after every sampling attempt (replenishment structure C), the probability of collecting a resource on any given attempt can be written as the probability that replenishment has occurred since the previous attempt. We refer to the time between consecutive attempts as the ‘first return time’ *w*, which follows a distribution *P*_*i*_ (*w*). We define 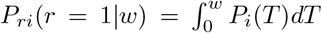 to be the probability that a replenishment r has occurred within a given duration *w*. We can then compute the average collection probability at an option *i* by averaging *P*_r*i*_(r = 1|*w*) over the distribution of first return times (Fig. 3e):

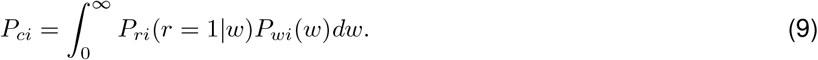

The distribution of first return times *P*_*wi*_(*w*) is solely a property of the policy. For example, if an agent can accurately track time and always returns to an option after a fixed duration, *P*_*wi*_(*w*) is a delta function; if an agent tries to return to an option after a fixed duration but has a noisy estimate of time, *P*_*wi*_(*w*) might be normally distributed. Here, since we have assumed that sampling attempts follow a Poisson process, the distribution of first return times is given by 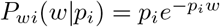.

In contrast, the replenishment probability depends solely on the statistics of replenishment through *P*_*i*_(T). As a result, the collection probability will generically depend on the form of *P*_*i*_(T). However, if all options again have the same form of replenishment statistics and only differ in their replenishment rates, we can write 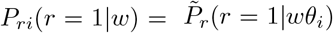. Again defining re-scaled variables 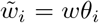 and 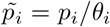, the collection probability can be written as:

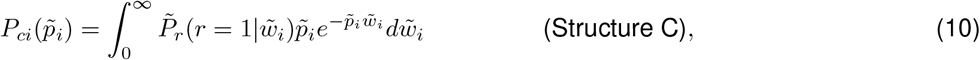

and the corresponding marginal gain as:

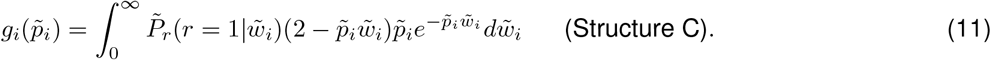

As before, because both *P*_*ci*_ and *g*_*i*_ depend only on 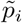, they are both equalized when 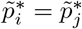 for all i, j (Fig. 3d).

In sum, when all options replenish independently of the agent’s actions, or when they all replenish after the agent samples an option (regardless of outcome), the optimal policy exhibits matching if all options share the same *form* of replenishment statistics, even if they differ in their replenishment *rates*.

### When optimality deviates from matching, relative collection probabilities depend only on the qualitative features of replenishment

In the previous section, we saw that the relationship between the marginal gain (which governs optimality) and the collection probability (which governs matching) differs depending on the replenishment structure. As a result, if individual options are governed by different replenishment structures, the optimal policy does not generally give rise to matching, even if the options share the same replenishment statistics (Fig. 4a; note the exception when replenishment follows a memoryless Poisson process). Even within a given replenishment structure, this relationship can depend on the *form* of replenishment (Eqs. 7-8 for structure B, and Eqs. 10-11 for structure C). As a result, if individual options are governed by the same replenishment structures but different forms of replenishment, the optimal policy does not generally give rise to matching (Fig. 4b; note the exception when all options are governed by replenishment structure A).

**Figure 4.**
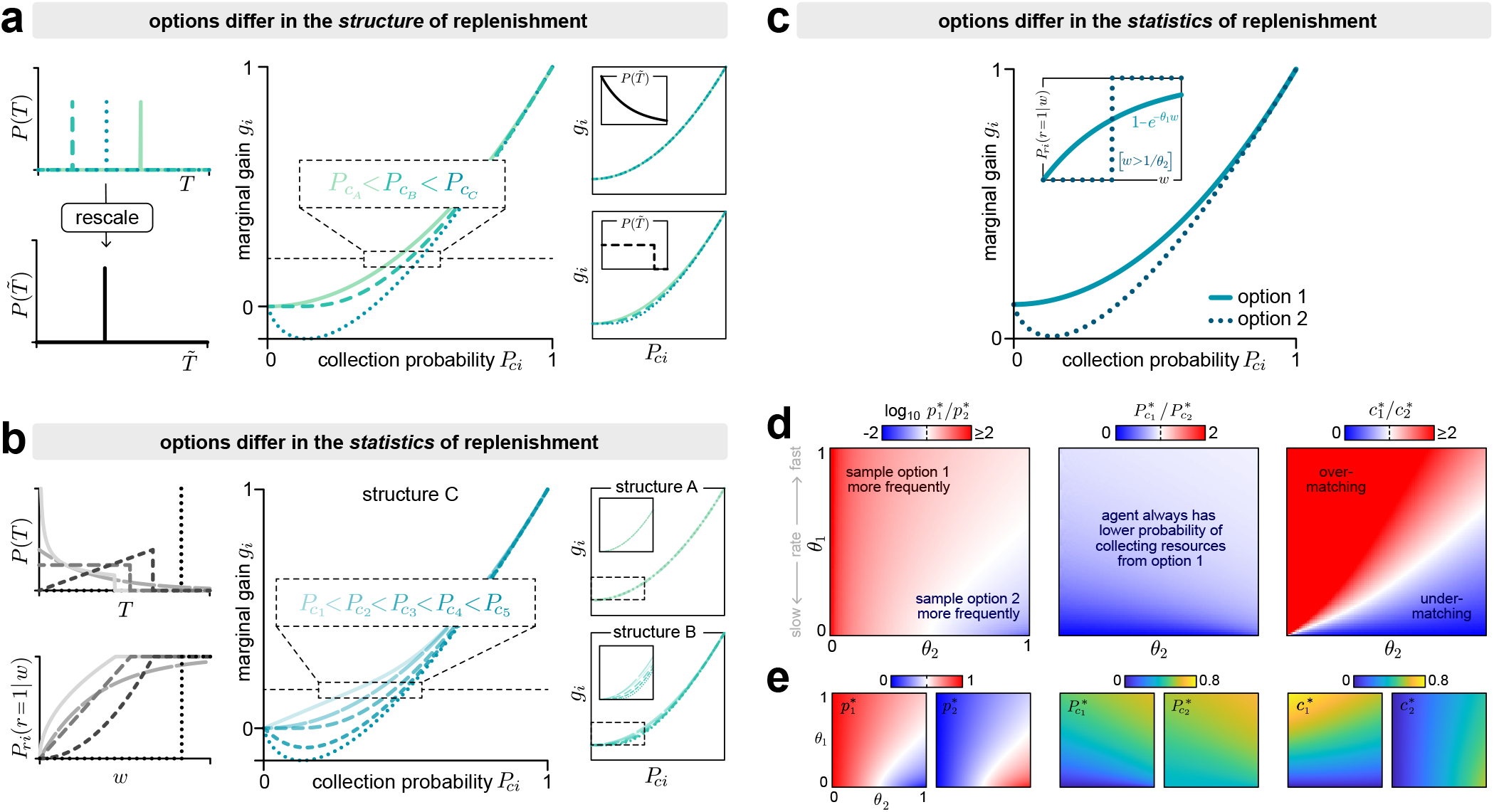
When the optimal policy deviates from matching, the rank-ordering of collection probabilities depends only on the qualitative features of replenishment. Optimality can deviate from matching when options differ in their replenishment structure but share the same statistics **(a)**, or when they share the same replenishment structure but differ in their statistics **(b-d). a)** To illustrate the effect of replenishment structure, we consider 3 options that are each governed by different replenishment structures but share the same replenishment statistics 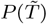, where 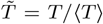 (left column). In general, the optimal policy does not necessarily give rise to matching, and the ordering of *P*_*c*_ depends on the replenishment structure (middle panel). The righthand panels show examples for other replenishment statistics; note that in the special case where the replenishment statistics follow a Poisson process (upper right), the optimal policy will always exhibit matching (with *P*_*ci*_ = *P*_*c*_). **b)** To illustrate the effect of replenishment statistics, we consider 5 options that are governed by the same replenishment structure but that differ in the form of their replenishment statistics *P* (*T*) and corresponding replenishment probabilities *P*_*r*_ (r = 1| *w*) (left column). Here again, the optimal policy does not always give rise to matching, and the ordering of *P*_*c*_ depends on the statistics of replenishment (middle panel). Since the optimal solution satisfies the condition that *g*_*i*_ is matched across options, the ordering of the collection probability under the optimal policy depends on the shape of *g*_*i*_(*P*_*ci*_), which in turn depends on both the structure and statistics of replenishment. When replenishment is governed by structure C (middle panel), the ordering of collection probabilities depends on the shape of *P*_*r*_(r = 1 |*w*) (with *P*_*c*1_ < *P*_*c*2_ < *P*_*c*3_ < *P*_*c*4_ < *P*_*c*5_). When replenishment is governed by structure B (lower right), the ordering depends on the shape of 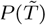 (with *P*_*c*2_ < *P*_*c*1_ < *P*_*c*3_ < *P*_*c*4_ < *P*_*c*5_). Note that in the special case where replenishment is governed by structure A (upper right), the optimal policy always exhibits matching (with *P*_*ci*_ = *P*_*c*_). **c)** To illustrate the effect of replenishment rates, we consider 2 options that share the same replenishment structure (structure C) but differ in the form and rates of their replenishment statistics. As in panel **b**, the optimal policy does not generically give rise to matching, and the ordering of collection probabilities depends on the shape of *P*_*r*_(r = 1 |*w*) (with *P*_*c*1_ < *P*_*c*2_). **d)** More generally, the optimal policy depends on the replenishment rates θ_1_ and θ_2_ (left panel; white denotes equal sampling rates for the two options). Under this policy, increasing the replenishment rate of either option leads to an increase in the collection probabilities of both options. However, the collection probability from option 1 is always lower than from option 2. In other words, the rank-ordering of collection probabilities is always preserved (middle panel; white denotes equal collection probabilities for the two options). The optimal policy therefore gives rise to under-matching when the collection rate from option 2 is higher (since in this case, the agent perceives option 2 to be more rewarding, but yet does not sample it enough to lower its collection probability to match that of option 1). When the collection rate from option 1 is higher, the optimal policy gives rise to over-matching (right panel; white denotes equal collection rates for the two options). **e)** Individual contributions of the ratios shown in panel **d**.

Furthermore, when optimality does not exhibit matching, the rank-ordering of collection probabilities depends only on these two qualitative features—the replenishment structure and the form of the replenishment statistics—and not on the values of the replenishment rates. To illustrate this point, we consider two options that are governed by the same replenishment structure but different replenishment statistics (Fig. 4c). Depending on their specific replenishment rates, the optimal policy involves sampling one or the other option more frequently (Fig. 4d-e, left panels). Under this policy, increasing the replenishment rate of either option will lead to an increase in the average collection probabilities of both options. Nevertheless, the same option always has the higher collection probability, regardless of which option is of higher quality (Fig. 4d-e, middle panels). As a result, the optimal policy can give rise to under-or over-matching depending on which option provides a higher collection rate (Fig. 4d-e, right panels), but the ordering of collection probabilities at optimality remains the same and can be predicted based on its relationship with the marginal gain (Fig. 4c).

### Low-quality environments produce larger deviations between optimality and matching

So far, we have explored the conditions under which the optimal policy does or does not exhibit matching. Here, we ask how much the optimal policy deviates from the matching policy that best maximizes the overall collection rate. This question is especially relevant because matching has been thought to be a feature of optimal or near-optimal behavior [4, 25], and simple neural network models for decision making have been proposed to account for the matching phenomenon [26].

As we have seen in the previous sections, any differences between optimality and matching fundamentally arise from differences in the relationship *g*_*i*_(*P*_*ci*_) between the marginal gain *g*_*i*_ and the collection probability *P*_*ci*_ across options. To better understand these differences, it is useful to consider general features of *g*_*i*_(*P*_*ci*_) that hold for all replenishment structures and statistics.

In the limit that the sampling rate goes to zero, the collection probability always goes to one (i.e., in the absence of any sampling, an option is guaranteed to be in the full state, and thus any sampling event is guaranteed to be successful). In the limit that the sampling rate becomes infinite, both the collection probability and its derivative always go to zero (i.e., under infinitely rapid sampling, an option is guaranteed to be in the empty state, and thus any sampling event is guaranteed to be unsuccessful). We can use these two sets of limits to bound the marginal gain:

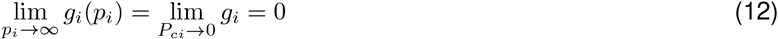

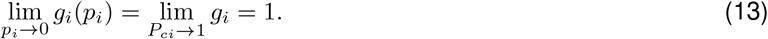

This can be seen by noting that g_*i*_ can be written as 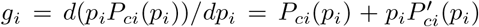, and taking the corresponding limits 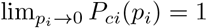 and 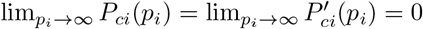.

The fact that the marginal gain always goes to one as the sampling rate goes to zero—regardless of the structure of statistics of replenishment—can be intuitively understood in terms of how the sampling rate p_*i*_ impacts the collection rate *c*_*i*_ = *p*_*i*_*P*_*ci*_(*p*_*i*_). An increase in sampling rate can impact the collection rate in two competing ways: (i) more frequent sampling leads to more opportunities for collecting a resource, thereby promoting an increase *c*_*i*_, and (ii) more frequent sampling reduces the probability that any given sampling event will lead to a collection, thereby promoting a decrease in *c*_*i*_. In the limit that the sampling rate is low, the first factor dominates the change in *c*_*i*_. In this limit, the average duration between sampling events is long, the resource is likely to be available when sampled, and thus any increase in sampling rate is likely to increase the collection rate. Thus,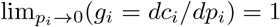.

Although the boundary values of *g*(*P*_*c*_) are the same for all environments, the derivatives of *g*(*P*_*c*_) depend on the details of the replenishment process. However, in the limit that the sampling rate goes to zero, the derivative of *g*(*P*_*c*_) does not depend on the structure nor the statistics of the replenishment process:

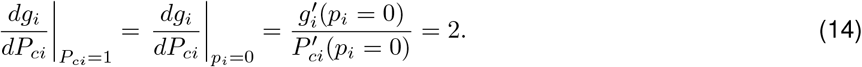

This can be seen by noting that 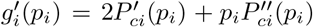, and taking the corresponding limit 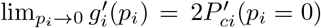.

Together, these constraints on the shape of *g*(*P*_*c*_) (Eqs. 13, 14) imply that optimal and matching policies become indistinguishable as the optimal collection probability goes to one, because the policy that equalizes the marginal gain will also equalize the collection probability (Fig. 5a). This limit is achieved when the environment is of sufficiently high quality and options replenish frequently. Indeed, even in settings that give rise to deviations between optimality and matching (Fig. 5b), these deviations tend to be small in high quality environments with high replenishment rates (Fig. 5c, upper right corners of heatmaps). In contrast, when the marginal gain at optimality is low, the collection probabilities across options can differ substantially, deviating markedly from matching (Fig. 5b). This can arise when at least one option is of low quality, such that the collection probability from that option is low even under optimal sampling (Fig. 5c, left panel; Fig. 5d). In such cases, the optimal and matching policies can also differ substantially (Fig. 5c, middle panel; Fig. 5e). When both options are of low quality, the overall collection rate is low under the optimal policy, but is much lower under the best matching policy (Fig. 5c, right panel; Fig. 5f).

**Figure 5.**
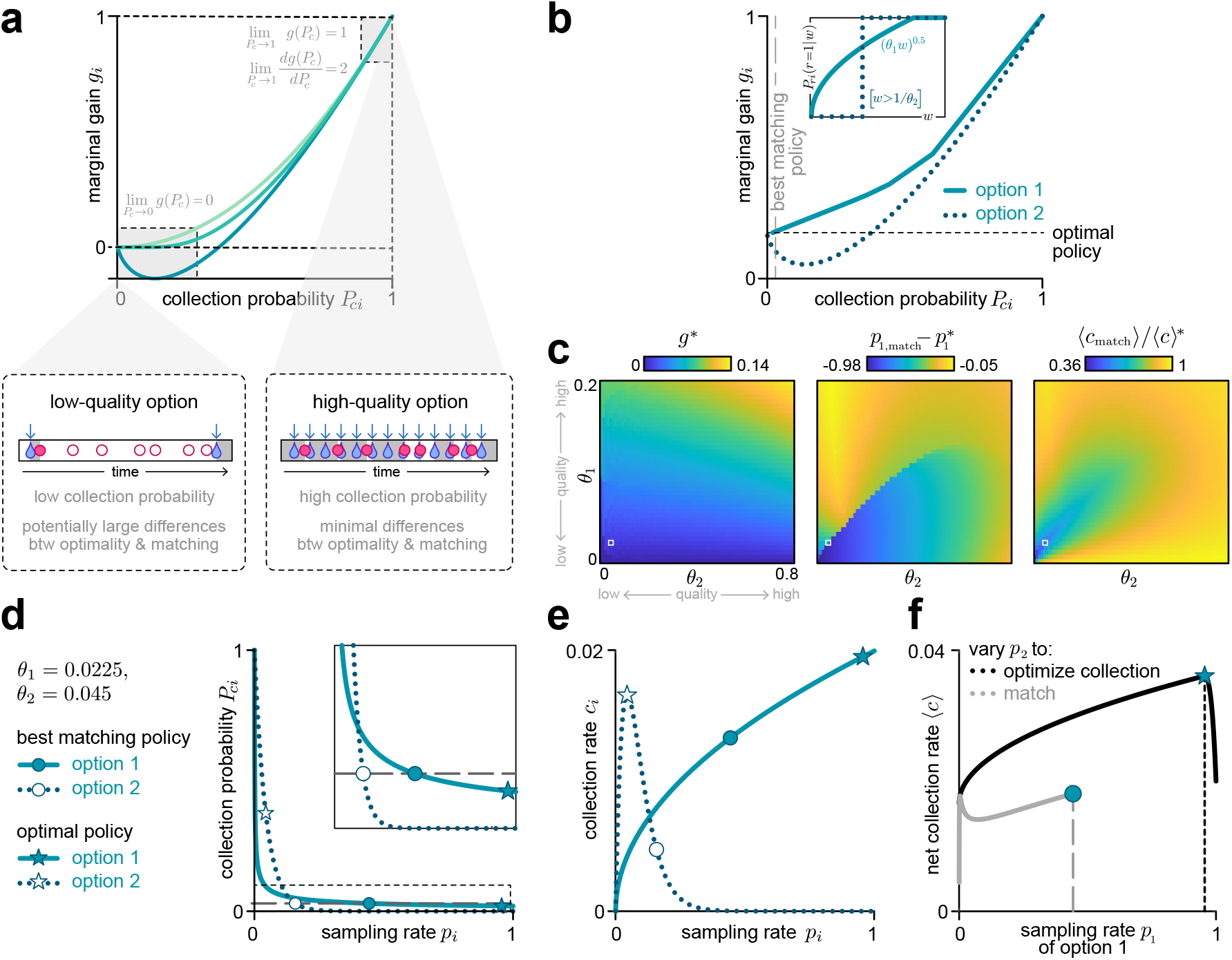
Low quality environments can lead to strong deviations between optimality and matching. **(a)** The marginal gain is guaranteed to go to zero as the collection probability goes to zero, regardless of the replenishment process. This situation can arise in low-quality environments where options replenish infrequently; in this regime, optimality can deviate significantly from matching (lower left). At the other extreme, as the collection probability goes to one, both the marginal gain and its slope have fixed limits regardless of the replenishment process. This situation can arise in high-quality environments where options replenish frequently; in this regime, optimality and matching are similar (lower right). **(b)** We consider an environment with two options that share the same replenishment structure (structure C) but differ in their replenishment statistics (inset). Due to the differences in replenishment statistics, the optimal policy (horizontal black dashed line) does not exhibit matching. The “best matching policy” that optimizes the net collection rate is marked by the vertical gray dashed line (main panel). **(c)** In low-quality environments where replenishment rates are low, the optimal and best matching policies can differ substantially from one another (lower left regions of heatmaps). In these regimes, the optimal policy has a low marginal gain (left), the sampling rates under the best-matching policy deviate from optimal sampling rates (middle), and the net collection rate under the best-matching policy is much lower than under the optimal policy (right). **(d-f)** We consider a specific example environment where θ_1_ = 0.0225 and θ_2_ = 0.045 (indicated by the white square markers in panel **c**). Circles and stars indicate the best matching and optimal policies, respectively. **(d)** Under the best matching policy, the collection probability is the same for both options (gray dashed horizontal line; see inset). Under the optimal policy, the collection probability differs substantially between the two options (stars). Inset shows expanded version of dashed box in main panel. **(e)** In this environment, the collection rate at option 2 changes non-monotonically with the sampling rate (dotted line). As a result, the optimal policy involves maximizing the collection rate at option 2 (open star), and allocating the remaining sampling rate budget to option 1 (filled star). Compared to the optimal policy, the best matching policy over-samples option 2 and under-samples option 1. **(f)** For any given value of the sampling rate *p*_1_, we compare the net collection rate ⟨*c*⟩ for an optimizing agent (i.e., an agent that adopts the value of *p*_2_ that maximizes the collection rate; black curve) with that of the matching agent (i.e., an agent that adopts the value of *p*_2_ such that the collection probabilities are equalized across options; gray curve). Note that both the optimizing and matching agents are subject to the same constraint on the total sampling rate; i.e., *p*_2_ <= 1 − *p*_1_. This constraint places a maximum limit on the value of *p*_1_ for the matching agent; above this value, matching cannot be achieved without *p*_2_ exceeding 1 − *p*_1_. The values of *p*_1_ for the optimal and best matching policies are indicated by the gray and black dashed vertical lines, respectively.

### Environmental fluctuations affect whether the optimal policy exhibits matching

So far, we have assumed that replenishment processes are stationary. However, in natural settings, the environment can fluctuate and can impact the replenishment process. We thus asked whether random fluctuations in the replenishment rates affect the relationship between optimality and matching.

We consider the scenario where all options share the same stationary replenishment structure and statistics, but where the replenishment rates can change at regular intervals. Whenever a change occurs, the replenishment rate θ_*i*_ of option *i* is drawn from a distribution *P*(θ_*i*_) with mean 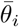 and variance 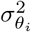. In this case, the conditions for optimality and matching are given by (see SI Section 1.1):

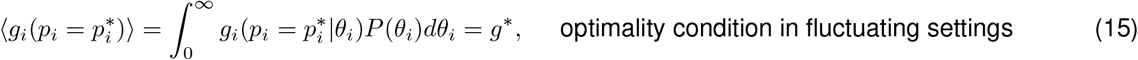

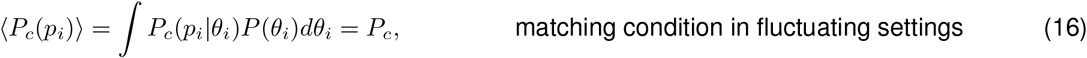

where the angular brackets ⟨·⟩ denote an average over the distribution of replenshment rates *P*(θ_*i*_), and where the marginal gain *g*_*i*_(*p*_*i*_|θ_*i*_) and collection probability *P*_*ci*_(*p*_*i*_|θ_*i*_) are defined as previously (Eqs. 1,2).

In previous sections, we showed that the marginal gain and collection probability can both be expressed as functions of a single variable *p*/θ, regardless of the replenishment structure (Eqs. 7-8, 10-11). This implies that if the distribution of replenishment rates *P*(θ) is controlled by two parameters that jointly specify the mean 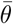 and variance 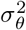 of the replenishment rates, the average marginal gain ⟨*g*⟩ and average collection probability ⟨*P*_*c*_⟩ will depend only on 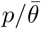 and 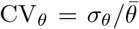 (see SI Section 2.1). Thus, when options are governed by the same replenishment structure and statistics but differ in their mean replenishment rates 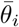, the optimal policy gives matching if all options fluctuate to the same degree (i.e., if 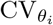 is matched across options). If options fluctuate by differing degrees, the optimal policy need not exhibit matching.

In such cases, the relative ordering of 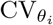 often determines the relative ordering of collection probabilities. For example, when options replenish after each collection (structure A) or when replenishment is governed by a Poisson process, the relationship between the marginal gain and collection probability is given by (from Eq. 5):

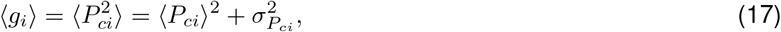

Where 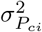 is the variance in the collection probability *P*_*ci*_ due to fluctuations in the corresponding replenishment rate θ_*i*_. Under these conditions, options with higher fluctuations—which in turn leads to higher variance in collection probabilities—will yield lower average collection probabilities under the optimal policy.

This finding holds more generally for other replenishment structures and statistics (see SI Section 2.2). To illustrate this, we consider two options that both replenish after each attempt, with replenishment times drawn uniformly (equivalently, we can consider options that replenish on regular intervals independently of the agent, with *P*(*T*|θ) = δ(*T*− θ)). We assume that the first option does not fluctuate 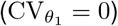, but the second option fluctuates uniformly over a range 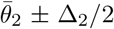 (such that 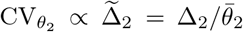 Fig. 6a). As these fluctuations increase, the optimal sampling rate of the fluctuating option can increase or decrease depending on the average replenishment rate 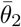 (Fig. 6b, left). However, the average collection probability is always lower from the fluctuating option compared to the reliable option (Fig. 6b, middle). As a result, larger fluctuations can lead to under-matching if the fluctuating option is perceived to be less rewarding (i.e., yields a lower collection rate), or over-matching if the fluctuating option is perceived to be more rewarding (i.e., yields a higher collection rate) (Fig. 6b, right).

**Figure 6.**
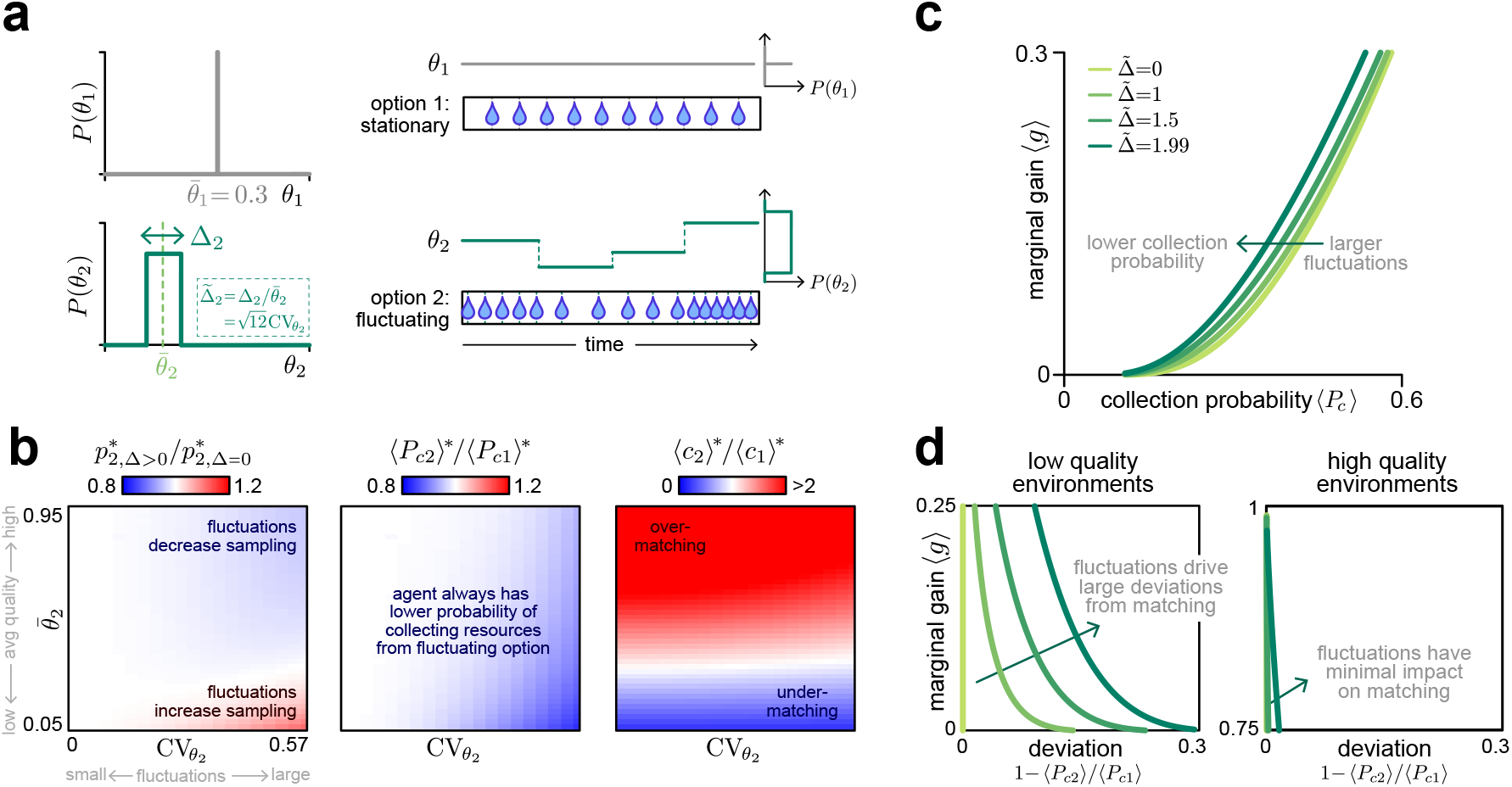
Environmental fluctuations impact whether the optimal policy exhibits matching. **(a)** We consider an environment with two options: the first option replenishes at a constant rate 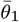, wh<u>ile</u> the second option replenishes at a rate that is drawn from a uniform distribution centered about 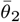 with width Δ_2_, such that 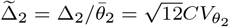. Both options replenish after each sample (structure C), with replenishment times drawn from a uniform distribution parameterized by 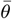 (this equivalently describes two options that replenish independently of the agent (structure B) with fixed replenishment times specified by 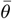, as schematized). For panels **b-d**, we set 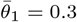. **(b)** The optimal sampling rate of the fluctuating option can decrease or increase with the degree of fluctuation, depending on the average replenishment rate (left). However, the relative probability of collecting resources from the second option always decreases as the fluctuations increase (middle). As a result, the optimal policy can either give rise to over-or under-matching, depending on whether the fluctuating option is perceived to be more or less rewarding (based on the relative collection rates) than the stationary option (right). **(c)** For a given marginal gain, the collection probability always decreases as fluctuations increase. **(d)** In low-quality environments where the marginal gain is low, fluctuations have a large impact on the relative collection probabilities (left panel). By contrast, in high quality environments where the marginal gain is high, the relative collection probabilities are only minimally affected by fluctuations (right panel).

To understand this result, we solve for the relationship between the average marginal gain and the average collection probability as a function of the degree of fluctuations 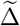 (Fig. 6c). For any fixed average marginal gain, larger fluctuations lead to a lower average collection probability. However, lower marginal gains—which arise in lower quality environments (Fig. 5a)—lead to more pronounced differences in collection probabilities between the two options (Fig. 6d, compare left and right panels). Together, this suggests that fluctuations drive deviations from matching, and these deviations are more pronounced in low-quality environments with low replenishment rates.

## Discussion

Since the first observation of the matching phenomenon in 1961, it has received widespread attention, and the degree of matching is now commonly reported in experiments where animals choose between multiple depleting options. However, despite the prevalence of these matching studies, the relationship between matching and optimality is not well understood. This is in part because matching is an emergent phenomenon that depends on how an animal’s choices interact with environmental dynamics to govern the availability of resources. As a result, it is often unclear how different experimental designs ‘should’ impact behavior, and whether one should be surprised by observations of matching. In this work, we address these questions by considering a large range of environments that differ in the structure and statistics of replenishment, and we ask whether matching is observed under the optimal policy for these different environments. Across these settings, we identify a range of conditions under which matching is optimal, and we isolate key environmental properties that govern deviations from matching.

Many experiments have been carried out with memoryless replenishment processes, where empty options are replenished at a fixed rate or probability per time, and where replenishment times are drawn from an exponential (or geometric, for tasks with discrete trials) distribution [2–5]. In such scenarios, the optimal fixed-sampling-rate policy is known to exhibit matching [2, 3, 22]. Nevertheless, our analysis shows that the optimal policy gives rise to matching across a range of other replenishment processes that vary in their structure, statistics, and rates, so long as the qualitative nature of the process is the same across options. If the options are instead governed by different types of replenishment processes, the optimal policy can deviate significantly from the matching policy, especially in low-quality environments where average replenishment rates are low. In such cases, the rank-ordering of collection probabilities depends only on the qualitative nature of the replenishment process, and not on the replenishment rates of different options. These findings provide testable predictions about the signatures of optimal behavior, and can serve as a guide for designing experiments that would yield large differences between optimality and matching.

Our characterization of optimality across different environments aligns with broader goals of understanding behavior in more naturalistic settings. Real environments not only consist of different food and water sources governed by different replenishment processes, but the rate at which these resources replenish can also be subject to weather and other environmental conditions. To better understand behavior in such scenarios, we explored how fluctuations in replenishment rates affect the relationship between optimality and matching. We found that the relative degree of fluctuation predicts relative differences in optimal collection probabilities—in particular, options with a higher degree of fluctuation always have a lower collection probability under the optimal policy, regardless of whether fluctuations preferentially impact low- versus high-quality options.

We interpreted these fluctuations as reflecting external variability in the environment itself, but they can equivalently be interpreted as reflecting internal uncertainty in the agent’s belief about the environment. More specifically, the optimal policy for a scenario in which replenishment rates are drawn from some distribution 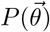 is the same as the optimal policy for a scenario where 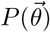 represents the agent’s belief about what the replenishment rates could be. This latter situation can arise during learning in a novel environment, when animals must discover the quality of different options through the outcomes of their actions. During early stages of learning, animals must make decisions based on the outcomes of sparse samples. In such settings, animals are likely to have a higher probability of collecting resources from higher-quality options, and are likely to bias their sampling toward those options [27]. This biased sampling would in turn lead to lower uncertainty in the estimated replenishment rates of high-quality options. Our results imply that carrying out the optimal policy under such uncertainties would continue to yield higher average collection probabilities for higher-quality options. This corresponds to under-matching, which has been observed in many previous experiments [14, 19, 21]. Furthermore, as animals update their beliefs and converge toward correct estimates of the true replenishment rates, relative differences in uncertainty between high- and low-quality options—and thus the degree of under-matching—would be expected to decrease. Indeed, this has been observed during early learning of a novel environment with multiple options [27], although in that setting, behavior was best described by a policy with memory of recent choice. Previous work has shown that under-matching can arise from learning over multiple time scales, which can be optimal in environments where reward rates regularly switch between two options [6]. Our work provides a complementary explanation for the ubiquity of under-matching: under-matching is a feature of optimality even in non-changing environments when the animal is more uncertain about the quality of the less rewarding options. Additional environmental dynamics—such as regular switches in reward rates—would further impact the agent’s belief about the underlying replenishment rates.

In this paper, we assumed that agents adopt a fixed-sampling-rate policy. Beyond enabling precise derivations of optimality and matching, such a policy is computationally compact and does not require the agent to remember its reward and choice histories. As such, it can potentially be implemented in simple neural networks [26]. However, real animals may adopt more complex behavioral strategies and may be subject to other behavioral constraints. For example, many animals have been observed to repeat their most recent choices, a phenomenon sometimes known as “stickiness” or “perseverance” [28–32], and logistic regression models fitted to behavioral data have often detected choice- and reward-history-dependent effects on animals’ next choices [27, 29–31]. Other experiments have shown that animals can track time [33, 34], which may allow them to exploit temporal regularities in the replenishment process. Our approach can potentially be adapted to explore the role of policy structure on optimality and matching, which remains an interesting question for future work.

## Acknowledgments

This work was motivated by multi-option foraging experiments that were conducted by Laura Grima and Josh Dudman [27], and we benefited from useful discussions with them about this work. YG and AMH were supported by the Howard Hughes Medical Institute. YG was also supported by the Agency for Science, Technology, and Research.

## Supplementary Information

## 1 Optimality condition

For a given fixed sampling rate policy 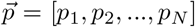 with Σ_i_ *p*_*i*_ ≤ 1, the overall average collection rate is given by: ⟨*c*⟩ = *c*_1_ +*c*_2_ +…+*c*_*N*_, where *c*_*i*_(*p*_*i*_) = *p*_*i*_*P*_*ci*_(*p*_*i*_) is the collection rate at an option *i* and *P*_*ci*_(*p*_*i*_) is the corresponding collection probability. Solving for the optimal policy 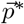 then corresponds to finding the sampling rates that maximizes ⟨*c*⟩ subject to the constraints that

For all the replenishment structures that we consider in this paper, 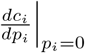 takes on the same value (of 1), and *c*_*i*_(*p*_*i*_) exhibits diminishing returns, i.e., 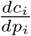 decreases with *p*_i_. This implies that if an agent is not sampling one of the options at all (e.g. *p*_j_ = 0 for option j) and sampling another option at a finite rate (e.g. *p*_k_ > 0 for option k), it is always possible to improve the overall average collection rate by decreasing the sampling rate at option k by a small amount δ*p* and instead sample option j at rate δ*p* (such that the total sampling rate is fixed). Formally, his is because the change in the average collection rate due to such a perturbation is given by: 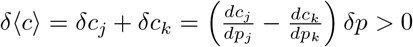.

This implies that the optimal sampling rate at each of the options must be strictly positive 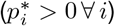. Therefore, the only potentially active constraint in our optimization problem is the one that involves the total sampling rate.

The optimal policy is therefore one that minimizes the following Lagrangian function:

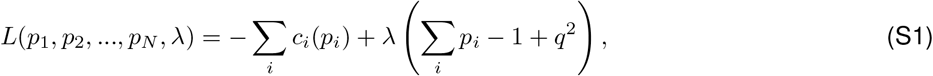

where we have introduced the slack variable *q* for the constraint.

The corresponding gradients of *L* with respect to its variables are then given by:

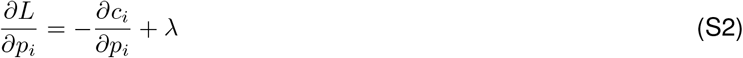

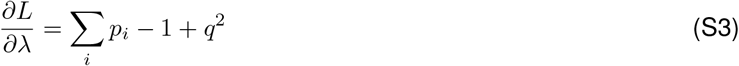

If the constraint is active (q = 0), the agent samples at the maximum possible rate (Σ_i_ *p*_i_ = 1and such that 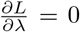), and the policy is optimal if:

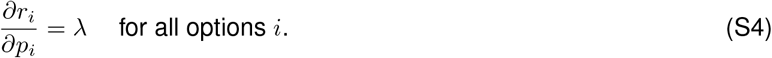

If the constraint is inactive (λ = 0 and q^2^ > 0), the agent samples less frequently that the maximum possible rate (Σ _i_ *p*_*i*_ < 1), and the policy is optimal if:

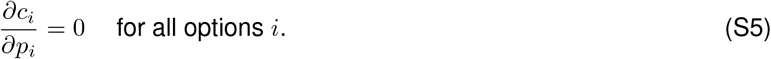

Therefore, in any given environment, the optimal policy 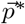 is one where 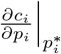 is the same across all options.

It is useful to note that when *c*_*i*_(*p*_*i*_) is a monotonically increasing function for at least one of the options in the environment, which is always the case for replenishment structures (A) and (B) regardless of the replenishment statistics, it is beneficial to sample as fast as possible and hence the constraint will be active (q = 0). In other words, in the space of replenishment processes we consider, an optimal policy with Σ_i_ *p*_*i*_ < 1 is only possible if all options have replenishment structure of type (C).

### 1.1 Optimality condition in the presence of replenishment rate fluctuations

If the replenishment rate θ_*i*_ of an option *i* is fixed, *P*_*c*_(*p*_*i*_|θ_*i*_) is the collection probability given a sampling rate *p*_*i*_ at that option. In the presence of replenishment rate fluctuations (in the form of changes in replenishment rates at regular intervals), if the values of θ_*i*_ are drawn from a distribution *P*(θ_*i*_), the corresponding average collection probability is then given by:

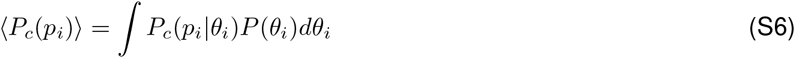

and the optimal policy is the one that maximizes the average collection rate ⟨*c*⟩ = ⟨*c*_1_⟩ + ⟨*c*_2_⟩ + … + ⟨*c*_*N*_ ⟩, with ⟨*c*_*i*_(*p*_*i*_)⟩ = *p*_*i*_⟨*P*_*c*_(*p*_*i*_)⟩.

Repeating the derivation for the optimality condition while taking into account the averaging over the distributions of replenishment rates, we find that the optimal policy 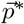 is one where 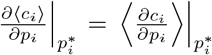 is the same across all options.

## 2 Relationship between the average marginal gain ⟨*g*⟩ and the average collection probability ⟨*P*_*c*_⟩ in the presence of fluctuations or uncertainty

### 2.1 For a given replenishment structure and statistics, the relationship between ⟨*g*⟩ and ⟨*P*_*c*_⟩ depends only on the coefficient of variation *CV* of *P* (*θ*)

We will assume that the distribution of replenishment rate *P*(θ) is a 2-parameter distribution, with both parameters jointly determining the mean 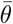 and standard deviation σ_θ_ of *P*(θ). The coefficient of variation is then defined to be 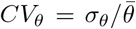. It is useful to define the scaled replenishment rate 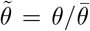, such that 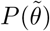 has mean 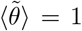 and standard deviation 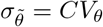.

For any function f(*p*/θ) that depends only on the ratio between the sampling rate and the replenishment rate, the average of the function over *P*(θ) is given by:

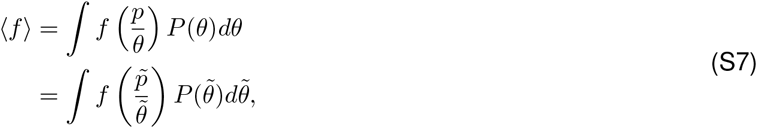

Where 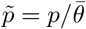.

Since 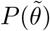 is a 1-parameter distribution with that parameter determined by *C*V_θ_, ⟨f⟩ is a function of both 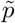 and *CV*_θ_. We saw in the main text that both the marginal gain *g* (Eqs. 5, 8, 11) and the collection probability *P*_*c*_ (Eqs. 4, 7, 10) are just functions of 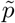. Therefore, if all options in the environment have the same replenishment structure and statistics but differ in their mean replenishment rates, even when their replenishment rates are not fixed, as long as they have the same degree of fluctuation *CV*_*θ*_, the relationship between ⟨*g*⟩ and ⟨*P*_*c*_⟩ is the same for all options, in which case the optimal policy gives rise to matching.

### 2.2 Increasing the degree of fluctuation *CV* for an option reduces the ratio of its collection probability to that of the other options under the optimal policy

We analyzed the relationship between ⟨*g*⟩ and ⟨*P*_*c*_⟩ for different replenishment structures and statistics, and for different distributions of replenishment rates *P*(θ). We considered both the uniform distribution and the gamma distribution for *P*(θ) (Fig. S1a) and found that in both cases, increasing *C*V of the distribution reduces ⟨*P*_*c*_⟩ for any given ⟨*g*⟩ that lies between 0 and 1 (Fig. S1b,c), This is the case across all the different environments and implies that under the optimal policy, ⟨*P*_*c*_⟩ is lower for options with a higher degree of fluctuation.

---

**Figure S1.**
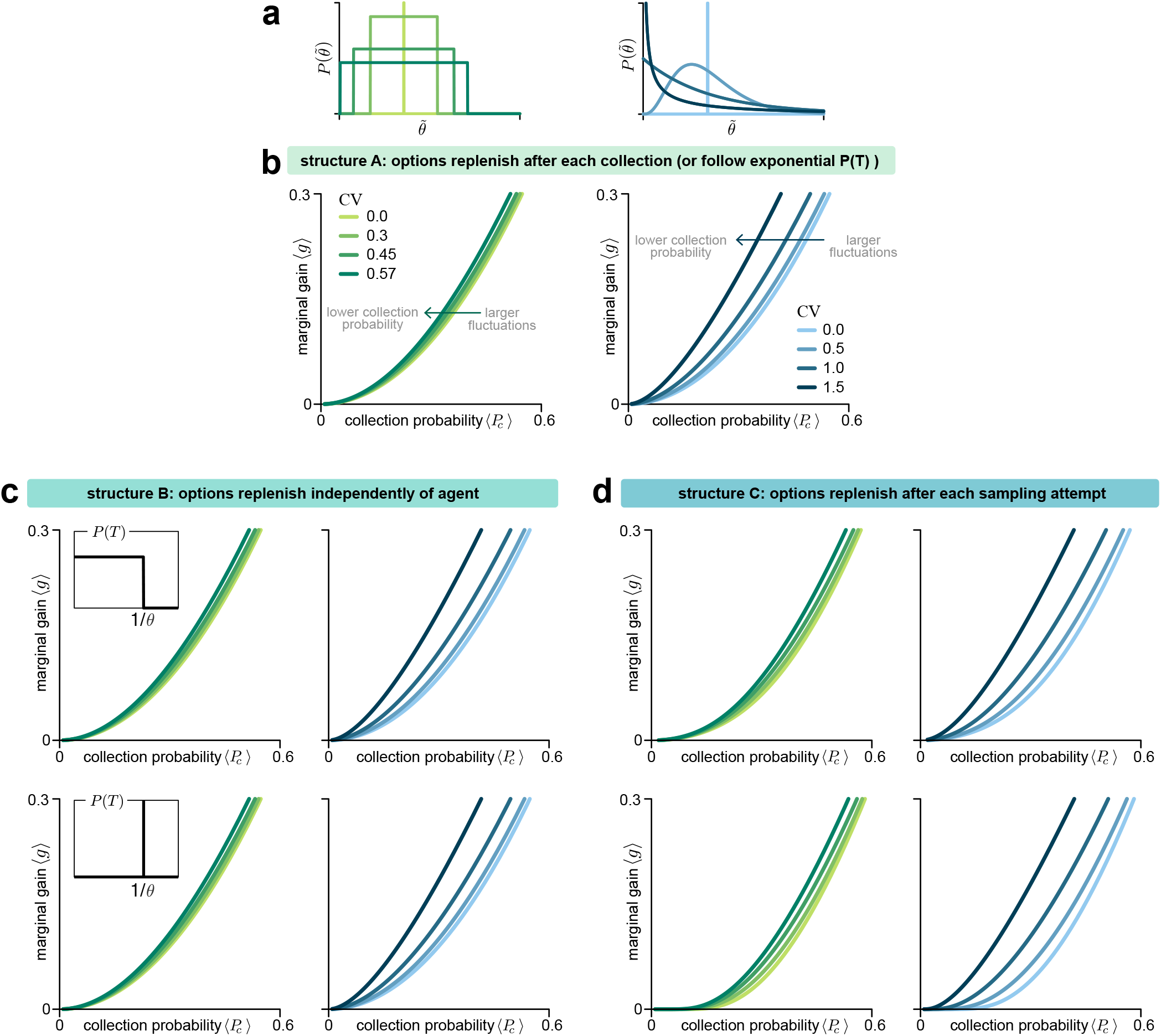
The impact of fluctuations on the relationship between marginal gain and collection probability is consistent across different environments. **(a)** We considered uniform distributions (left) and gamma distributions (right) for the distribution of replenishment rates *P* (θ). **(b)** The relationship between ⟨*g*⟩ and ⟨*P*_*c*_⟩ when the replenishment process is of structure type (A), or if *P* (T) is exponential. **(c-d)** The relationship between ⟨*g*⟩ and ⟨*P*_*c*_⟩ for other replenishment structures and statistics (top row: *P* (T) is uniform; bottom row: *P* (T) is a delta-distribution.)

